# Genetic background influences cocaine neuroplasticity in the accumbens shell of male mice

**DOI:** 10.64898/2026.09.21.753182

**Authors:** Chinonso A. Nwakama, Andrew D. Chapp

**Affiliations:** Medical Scientist Training Program, Icahn School of Medicine at Mount Sinai, New York, NY 10029; Nash Family Department of Neuroscience, Friedman Brain Institute, Icahn School of Medicine at Mount Sinai, New York, NY 10029; Department of Neuroscience, Feinberg School of Medicine, Northwestern University, Chicago, IL 60611; Department of Neuroscience, University of Minnesota, Minneapolis, MN 55455

## Abstract

Genetics shape cocaine behavior and abuse liability, but the underlying neurophysiological effects remain understudied. In male C57BL/6J, B6129SF2/J, and DO/J mice, we report conserved and divergent cocaine-induced neuroplasticity in the accumbens shell following 5 days of cocaine and 10–14 days of abstinence.

## INTRODUCTION

Globally, cocaine use disorder (CUD) remains an important contributor to the burden of disease(*1*). In rodents and humans, genetic background has been shown to drastically alter the behavioral effects of cocaine. In rodents, this includes cocaine locomotor sensitization(*2, 3*) and cocaine self-administration(*4*). In parallel, candidate gene association and genome wide association studies have attempted to identify genetic markers which may influence susceptibility to cocaine use disorder(*5, 6*). While these approaches have shed light on behavioral differences and genetic predispositions for cocaine sensitivity, they have yet to show how these differences in genetics may shape cocaine neuroplasticity.

Cocaine-induced neuroplastic changes in the brain’s reward circuit, such as the nucleus accumbens shell (NAcSh), are thought to contribute to CUD(*7, 8*). Comparing and contrasting these effects across genetically distinct rodent populations is an important step towards understanding how genetic background may contribute to individual susceptibility to cocaine dependence. As such, we investigated whether genetic background influences cocaine-induced neuroplasticity in the NAcSh of three populations of male mice with increasing genetic heterogeneity: C57BL/6J, B6129SF2/J, and Diversity Outbred/J (DO/J). Notably, we have previously reported differences in cocaine locomotor sensitization in these three strains of male mice [C57BL/6J > B6129SF2/J = DO/J](*3*). Recording from 150 medium spiny neurons in the medial portion of the NAcSh after 5 daily injections of cocaine and 10-14 days of abstinence, we report both conserved and divergent cocaine-induced neuroplasticity among these three strains. Our findings highlight a genetic diversity in the neuropharmacological actions of cocaine on abstinence plasticity in the accumbens shell of male mice.

## MATERIAL AND METHODS

Animal procedures were performed at the University of Minnesota in facilities accredited by the Association for Assessment and Accreditation of Laboratory Animal Care (AAALAC) and in accordance with protocols approved by the University of Minnesota Institutional Animal Care and Use Committee (IACUC), as well as the principles outlined in the National Institutes of Health Guide for the Care and Use of Laboratory Animals. Male C57BL/6J (#000664), B6129SF2/J (#101045), and DO/J (#009376) mice were obtained from The Jackson Laboratory (Bar Harbor, ME). Detailed information on experimental design, electrophysiology recording and statistics can be found in the Supplemental Materials.

## RESULTS

### Cocaine-induced reduction in excitability within NAcSh MSNs is conserved across strains

Cocaine-induced neuroadaptations within the nucleus accumbens are thought to contribute to relapse behavior(*7–9*), we focused our effort on comparing and contrasting the ramifications of 10-14 days abstinence on medium spiny neuron neurophysiology in the NAcSh among these three strains. All three strains of mice displayed a conserved reduction in medium spiny neuron excitability following 5 daily injections of cocaine (Fig 1A) and a 10-14 day abstinence period prior to recordings (C57BL/6J: two-way ANOVA-RM, F(_8,216_)=10.69, interaction, p<0.0001, B6129SF2/J: two-way ANOVA-RM, F(_8,216_)=6.38, interaction, p<0.0001, DO/J: two-way ANOVA-RM, F(_8,168_)=4.52, interaction, p<0.0001, Fig 1B-D).

**Fig 1.**
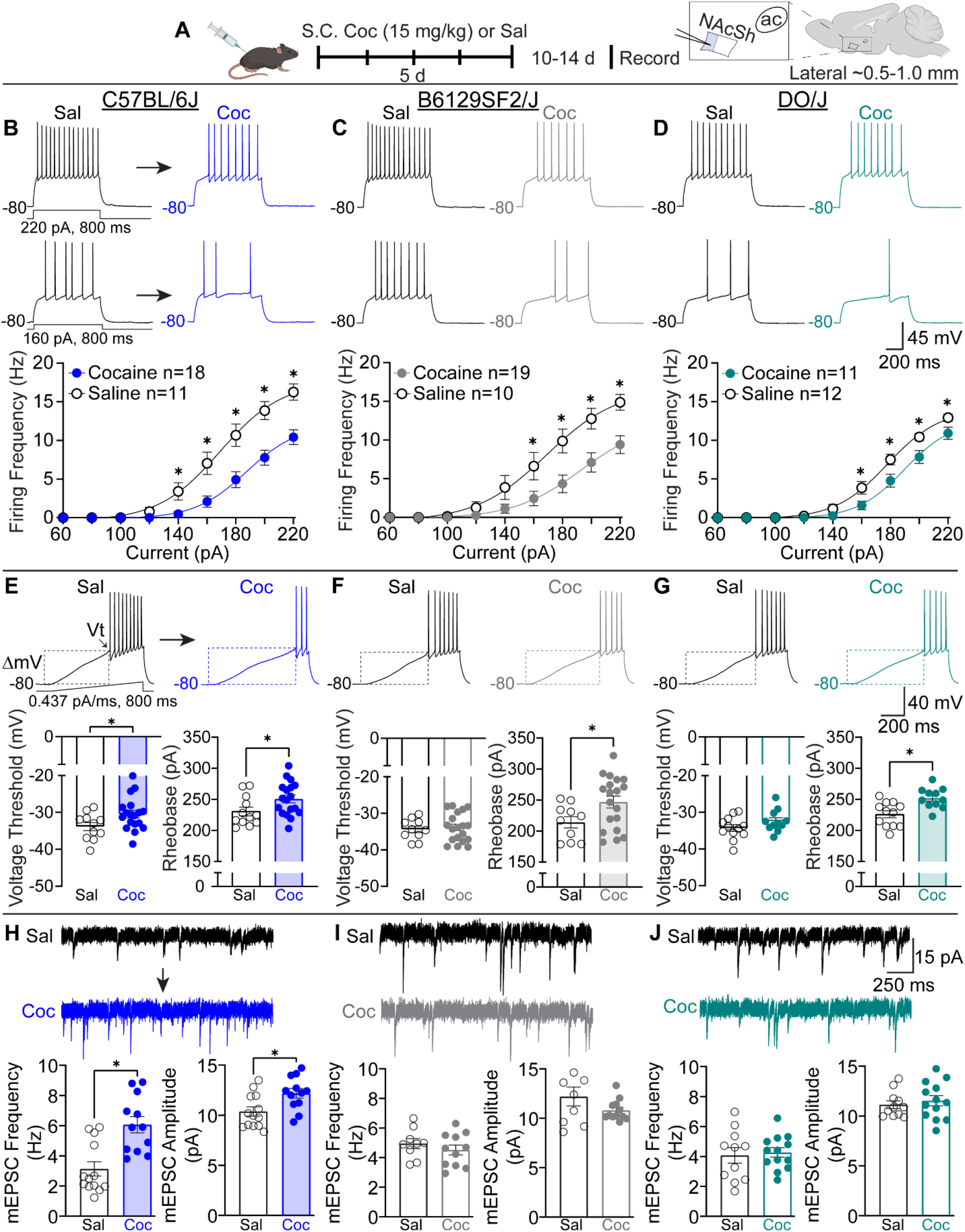
Conserved and divergent cocaine neuroplasticity in male NAcSh MSNs. **A** Experimental timeline and recording map. **B** Top: Representative excitability traces for saline (black) and cocaine (blue) at +160 and +220 pA in C57BL/6J male mice. Bottom: Summary current injection response curve for saline (black) and cocaine (blue) in C57BL/6J male mice, *p<0.05. **C** Top: Representative excitability traces for saline (black) and cocaine (gray) at +160 and +220 pA in B6129SF2/J male mice. Bottom: Summary current injection response curve for saline (black) and cocaine (gray) in B6129SF2/J male mice, *p<0.05. **D** Top: Representative excitability traces for saline (black) and cocaine (cyan) at +160 and +220 pA in DO/J male mice. Bottom: Summary current injection response curve for saline (black) and cocaine (cyan) in DO/J male mice, *p<0.05. **E** Top: Representative ramp trace for saline (black) and cocaine (blue) in C57BL/6J male mice. Bottom: Summary data for voltage threshold to action potential (left) and rheobase (right), *p<0.05. **F** Top: Representative ramp trace for saline (black) and cocaine (gray) in B6129SF2/J male mice. Bottom: Summary data for voltage threshold to action potential (left) and rheobase (right), *p<0.05. **G** Top: Representative ramp trace for saline (black) and cocaine (cyan) in DO/J male mice. Bottom: Summary data for voltage threshold to action potential (left) and rheobase (right), *p<0.05. **H** Top: Representative mEPSC traces for saline (black) and cocaine (blue) in C57BL/6J mice. Bottom: Summary data for mEPSC frequency (left) and mEPSC amplitude (right), *p<0.05. **I** Top: Representative mEPSC traces for saline (black) and cocaine (gray) in B6129SF2/J mice. Bottom: Summary data for mEPSC frequency (left) and mEPSC amplitude (right). **J** Top: Representative mEPSC traces for saline (black) and cocaine (cyan) in DO/J mice. Bottom: Summary data for mEPSC frequency (left) and mEPSC amplitude (right). Excitability studies contained n=3-4 mice per treatment and glutamatergic studies contained n=3-4 mice per treatment group. Abbreviations: Saline (sal), cocaine (coc), and subcutaneous (S.C.).

### Cocaine-induced alteration to voltage threshold is C57BL/6J specific while effects on rheobase are conserved across strains

Cocaine exposure and abstinence have been shown to alter voltage gated sodium channel properties(*10*), and we have reported this finding previously in C57BL/6J where cocaine-induced a depolarizing shift in voltage threshold to elicit an action potential(*11*). Here, we recapitulate that finding and demonstrate that it is C57BL/6J specific (C57BL/6J: Welch’s t-test, p=0.036, B6129SF2/J: Welch’s t-test, p=0.71, and DO/J: Welch’s t-test, p=0.15, Fig 1E-G). Conversely, the amount of current required to elicit an action potential (rheobase) was significantly increased in all strains (C57BL/6J: Welch’s t-test, p=0.041, B6129SF2/J: Welch’s t-test, p=0.022, DO/J: Welch’s t-test, p=0.002, Fig 1E-G).

### Cocaine-induced alteration to glutamatergic strength is C57BL/6J specific

Cocaine-induced alterations to the glutamatergic system in the accumbens has been postulated as a contributing factor in relapse(*9, 12, 13*). Part of these glutamatergic alterations include an increase in mEPSC frequency and amplitude following cocaine exposure and 10-14 days of abstinence(*11, 14*). This alteration has been previously reported in male C57BL/6J mice(*11, 12*). We recapitulate this finding and show that it is not conserved across mouse strains (Fig 1H-J). Specifically, mEPSC frequency was only increased in male C57BL/6Js (C57BL/6J: Welch’s t-test, p=0.0005, B6129SF2/J: Welch’s t-test, p=0.37, DO/J: Welch’s t-test, p=0.76) and mEPSC amplitude was only increased in male C57BL/6Js (C57BL/6J: Welch’s t-test, p=0.012, B6129SF2/J: Welch’s t-test, p=0.19, DO/J: Welch’s t-test, p=0.54).

### No major differences in passive membrane properties among mouse strains

Resting membrane potential (C57BL/6J Sal: −73.6 ± 1.04; C57BL/6J Coc: −75.3 ± 1.05; B6129SF2/J Sal: −75.5 ± 1.8; B6129SF2/J Coc: −77.3 ± 1.0; DO/J Sal: −75.2 ± 1.04 and DO/J Coc: −76.1 ± 1.2, one-way ANOVA, F(_5,75_) = 1.11, p=0.36) and membrane capacitance (C57BL/6J Sal: 101.6 ± 6.8; C57BL/6J Coc: 91.2 ± 3.6; B6129SF2/J Sal: 102.0 ± 11.9; B6129SF2/J Coc: 88.2 ± 2.5; DO/J Sal: 105.0 ± 7.0; and DO/J Coc: 103.0 ± 7.8, one-way ANOVA, F(_5,75_) = 1.16, p=0.17) were not significantly different. Membrane resistance was significantly different (C57BL/6J Sal: 79.4 ± 5.0; C57BL/6J Coc: 82.4 ± 2.4; B6129SF2/J Sal: 85.7 ± 4.4; B6129SF2/J Coc: 82.2 ± 3.7; DO/J Sal: 96 ± 4.2, and DO/J Coc: 92.4 ± 4.6, one-way ANOVA, F(_5,75_) = 2.51, p=0.04), however multiple pairwise comparisons revealed no significant differences between groups.

## DISCUSSION

Using the same mice strains, where we previously reported significant differences in cocaine locomotor sensitization(*3*), we now show that these three strains have both conserved and divergent cocaine-induced neuroplastic changes in the NAcSh following abstinence. Conserved cocaine-induced neuroplasticity includes a reduction in medium spiny neuron excitability and increased current required to elicit an action potential. Divergent cocaine-induced neuroplasticity includes a depolarizing shift in voltage threshold to elicit an action potential, and cocaine-induced potentiation in glutamatergic strength which are only observed in the C57BL/6J mice. These differences in neurophysiological findings appear to correlate with behavioral sensitization, in that the C57BL/6J male mice had a greater cocaine locomotor response compared to the B6129SF2/J and the DO/J male mice(*3*).

Previous work within male C57BL/6J mice has demonstrated that potentiation and manipulation of glutamatergic strength in the NAcSh during drug abstinence can alter relapse(*12*). The B6129SF2/J and DO/J mice do not share this neurophysiological homology during cocaine abstinence. Of the individuals who experiment with drugs such as cocaine, only a subset progress to dependence. This then begs the question of whether genetic variability in cocaine use and progression to dependence in humans can be better modeled in genetically diverse rodent strains. Is the male C57BL/6J a “good” mouse model for CUD in humans, or is it the exception to the rule? Key questions remain. If neurophysiology predicts drug sensitivity or relapse, or is used as a defining characteristic in these behavioral measures, should therapeutic treatment strategies target neurophysiology that is different and correlates with enhanced drug sensitivity (e.g., glutamatergic aspect observed in C57BL/6J males), or should treatment strategies focus on properties conserved across strains and species (e.g., decreased medium spiny neuron excitability)?

Boudreau *et al*, reported increased AMPAR expression in the accumbens of Sprague Dawley rats was associated with cocaine behavioral sensitization(*15*). These AMPAR adaptations were not present 1 day after the last cocaine exposure but were present after 21 days of abstinence. Meanwhile, Zhang *et al*, reported in the same strain of rats that cocaine precipitated alterations to whole-cell sodium currents in medium spiny neurons of the accumbens. These alterations were detectable 3 days after the last cocaine exposure, although the authors did not record in later withdrawal periods(*10*). The C57BL/6J mice display both these cocaine-induced neuroadaptations in the 10-14 day withdrawal window. Do glutamatergic plasticity and alterations to voltage gated sodium channels occur concurrently, or does one form of plasticity drive compensations to the other? The lack of glutamatergic plasticity in the B6129SF2/J and DO/J paired with a lack of plasticity in voltage gated sodium channels highlights that both forms of cocaine-induced neuroplasticity are potentially tightly correlated.

We acknowledge that a limitation of this study was that electrophysiology recordings were conducted in male mice only. Given that we have reported divergent behavioral and electrophysiology responses in the C57BL/6J strain based on sex and estrous cycle(*11*), future studies should examine the effects of cocaine exposure and abstinence in females among these strains to determine potential sex and strain specific alterations in cocaine-induced neuroplasticity in the NAcSh. Having a full behavioral and neurophysiological profile among strains by sex for drugs of abuse may aid in the development of therapeutic targets for treatment in humans.

## Acknowledgements

We would like to thank Dr. Andréa Collins, Dr. Timothy W Chapp, Dr. Hannah M. McMullan, and Dr. Qing-Hui Chen for their proof reading, as well as Dr. Paul G. Mermelstein and Dr. Mark J. Thomas for obtaining funding. A version of this manuscript has been deposited on biorxiv.

## Author Contributions

A.D.C. and C.A.N designed and performed experiments; A.D.C. and C.A.N. analyzed data; A.D.C. and C.A.N. prepared figures; A.D.C. and C.A.N. drafted manuscript; A.D.C. and C.A.N. interpreted results of experiment; A.D.C. and C.A.N. edited and revised manuscript.

## SUPPLEMENTAL MATERIALS AND METHODS

### Animal Housing

C57BL/6J and B6129SF2/J were group housed, while DO/J mice were individually housed due to a history of aggressive behavior(*1*). All animals were kept on a 10:14 light:dark cycle with food and water ad libitum(*2, 3*).

### Chemicals

All chemicals were obtained from Sigma-Aldrich (St Louis, MO, USA), except cocaine hydrochloride and isoflurane (Boynton Pharmacy, University of Minnesota, MN, USA).

### Cocaine and saline injections

Mice were randomly divided into groups and received either a subcutaneous saline or cocaine (15 mg/kg) injection daily for 5 days.

### Whole-cell recordings

Mice (8-14 weeks old) in cocaine abstinence (10-14 days after the last behavioral day) were used for electrophysiology recordings. Animals were sacrificed between 09:00 and 17:00. Mice were anesthetized with isoflurane (3% in O_2_) and decapitated. The brain was rapidly removed and chilled in ice cold cutting solution, containing (in mM): 228 sucrose, 2.5 KCl, 7 MgSO_4_, 1.0 NaH_2_PO_4_, 26 NaHCO_3_, 0.5 CaCl_2_, 11 d-glucose, pH 7.3-7.4, continuously gassed with 95:5 O_2_:CO_2_ to maintain pH and pO_2_. A brain block was cut including the NAcSh region and affixed to a vibrating microtome (Leica VT 1000S; Leica, Nussloch, Germany). Sagittal sections of 240 µm thickness were cut, and the slices transferred to a holding container of artificial cerebral spinal fluid (ACSF) maintained at 30 °C, continuously gassed with 95:5 O_2_:CO_2_, containing (in mM): 119 NaCl, 2.5 KCl, 1.3 MgSO_4_, 1.0 NaH_2_PO_4_, 26.2 NaHCO_3_, 2.5 CaCl_2_, 11 d-glucose, and 1.0 ascorbic acid (osmolality: 295–302 mOsmol L^−1^; pH 7.3-7.4)(*4, 5*) and allowed to recover for 1 hr. Following recovery, slices were transferred to a glass-bottomed recording chamber and viewed through an upright microscope (Olympus) equipped with DIC optics, and an IR-sensitive video camera (DAGE-MTI).

Slices transferred to the glass-bottomed recording chamber were continuously perfused with ACSF, gassed with 95:5 O_2_:CO_2_, maintained at room temperature and circulated at a flow of 2 mL min^-1^. Patch electrodes were pulled (Flaming/Brown P-97, Sutter Instrument, Novato, CA) from borosilicate glass capillaries with a tip resistance of 5–10 MΩ. Electrodes were filled with a solution containing (in mM) 135 K-gluconate, 10 HEPES, 0.1 EGTA, 1.0 MgCl_2_, 1.0 NaCl, 2.0 Na_2_ATP, and 0.5 Na_2_GTP (osmolality: 280–285 mOsmol L^−1^; pH 7.3)(*6*). MSNs were identified under IR-DIC based on morphology and their hyperpolarizing membrane potential (−70 to −80 mV) and were voltage clamped at −80 mV using a Multiclamp 700B amplifier (Molecular Devices), currents filtered at 2 kHz and digitized at 10 kHz. Holding potentials were not corrected for the liquid junction potential. Once a GΩ seal was obtained, slight suction was applied to break into whole-cell configuration and the cell was allowed to stabilize which was determined by monitoring capacitance, membrane resistance, access resistance and resting membrane potential (V_m_)(*6–8*). Cells that met the following criteria were included in the analysis: action potential amplitude ≥50 mV from threshold to peak, resting *V*_m_ negative to −64 mV, and <20% change in series resistance during the recording. Passive membrane properties, capacitance and membrane resistance were measured from the membrane test in pClamp (Molecular Devices). Resting neuronal membrane potential reported in Table S1 was recorded immediately after breaking into whole-cell mode(*2, 6, 9–11*).

To measure NAcSh MSN neuronal excitability, V_m_ was adjusted to −80 mV by continuous negative current injection, and a series of square-wave current injections was delivered in steps of +20 pA, each for a duration of 800 ms. To determine the action potential voltage threshold (Vt), and rheobase, ramp current injections (0.437 pA/ms, 800 ms) were made from a holding potential of −80 mV.

For miniature excitatory postsynaptic current recordings (mEPSC)(*2*), slices transferred to the glass-bottomed recording chamber were continuously perfused with ACSF containing lidocaine (0.7 mM) to block voltage-gated sodium channels and picrotoxin (100 μM) to block GABAR, and was continuously gassed with 95:5 O_2_:CO_2_, maintained at room temperature and circulated at a flow of 2 mL min^-1^. Patch electrodes were pulled from borosilicate glass capillaries with a tip resistance of 5–10 MΩ and whole-cell recordings were made. Electrodes were filled with a cesium methanesulfonate (CsMeSO_4_) solution containing (in mM): 120 CsMeSO4, 15 CsCl, 10 TEA-Cl, 10 HEPES, 0.4 EGTA, 8.0 NaCl, 2.0 Na2ATP, and 0.3 Na2GTP (osmolality: 280–285 mOsmol L−1; pH 7.3). MSNs were identified under IR-DIC based on morphology and their hyperpolarizing membrane potential (−70 to −80 mV) and were voltage clamped at −80 mV using A Multiclamp 700B amplifier (Molecular Devices), currents filtered at 2 kHz and digitized at 10 kHz. Holding potentials were not corrected for the liquid junction potential. mEPSCs were recorded for 2 minutes and analyzed offline using Mini Analysis software (synaptosoft) with an amplitude threshold set at three times the noise level.

### Statistical analysis

Data values were reported as mean ± SEM. All statistical analyses were performed with a commercially available statistical package (GraphPad Prism, version 9.4.1). Probabilities less than 5% were deemed significant *a priori*. Depending on the experiments, group means were compared using an unpaired Welch’s *t*-test, a one-way ANOVA, or a two-way ANOVA repeated measures. Where differences were found Holm-Sidak post-hoc tests were used for multiple pair-wise comparisons.

